# Genome-wide repressive capacity of promoter DNA methylation is revealed through epigenomic manipulation

**DOI:** 10.1101/381145

**Authors:** Keegan Korthauer, Rafael A. Irizarry

## Abstract

The scientific community is increasingly embracing open science. This growing commitment to open science should be applauded and encouraged, especially when it occurs voluntarily and prior to peer review. Thanks to other researchers’ dedication to open science, we have had the privilege of conducting a reanalysis of a landmark experiment published as a preprint with data made available in a public repository. The study in question found that promoter DNA methylation is frequently insufficient to induce transcriptional repression, which appears to contradict a large body of observational studies showing a strong association between DNA methylation and gene expression. This study was the first to evaluate whether forcibly methylating thousands of DNA promoter regions is sufficient to suppress gene expression. The authors’ data analysis did not find a strong relationship between promoter methylation and transcriptional repression. However, their analyses did not make full use of statistical inference and applied a normalization technique that removes global differences that are representative of the actual biological system. Here we reanalyze the data with an approach that includes statistical inference of differentially methylated regions, as well as a normalization technique that accounts for global expression differences. We find that forced DNA methylation of thousands of promoters overwhelmingly represses gene expression. In addition, we show that complementary epigenetic marks of active transcription are reduced as a result of DNA methylation. Finally, by studying whether these associations are sensitive to the CG density of promoters, we find no substantial differences in the association between promoters with and without a CG island. The code needed to reproduce are analysis is included in the public GitHub repository github.com/kdkorthauer/repressivecapacity.

## Introduction

The basic biology and biomedical communities are rapidly evolving as they increasingly embrace open science. Providing rapid and open access to results and data improves research efficiency and the robustness of scientific findings (Munafò et al., 2017). This growing commitment to open science should be applauded and encouraged, especially when it occurs voluntarily and prior to peer review. Thanks to other researchers’ dedication to open science, we have had the privilege of conducting a reanalysis of a landmark experiment published as a preprint (Ford et al., 2017) with data made available in a public repository.

The recent study by Ford et al. (2017) posited that promoter DNA methylation is frequently insufficient to induce transcriptional repression. This finding is unexpected given the large body of research that demonstrates the a relationship between DNA methylation and gene expression. Methylation of CG islands within promoters has been shown to be strongly negatively correlated with gene expression (Jones, 2012; Schübeler, 2015). While it is generally accepted that DNA methylation plays a prominent role in gene regulation, there is debate over whether it acts as a causal signal or occurs as a downstream effect.

Although it has not been shown systematically that DNA methylation acts as a primary and causal signal genome-wide, it has been shown to be sufficient for transcriptional repression of individual genes (Razin and Cedar, 1991; Jackson-Grusby et al., 2001; Yang et al., 2012; Bintu et al., 2016; Amabile et al., 2016; Kungulovski et al., 2015), and even necessary for some mechanisms of long term promoter silencing such as X-inactivation and imprinting (Li et al., 1993; Jaenisch and Bird, 2003; Bestor et al., 2015). Transcriptional silencing is also stimulated by a complex network of epigenetic marks, including methylation and deacetylation of certain histone subunits of nucleosomes near the transcription start site (Kungulovski et al., 2015), which are thought to be regulated in part by the recruitment of methylated DNA-binding proteins (Klose and Bird, 2006). It is unknown whether the primary role of DNA methylation in this complex network is initiation or maintenance of repressive signals.

The recent study by Ford et al. (2017) was the first to comprehensively evaluate whether the addition of DNA methylation to endogenous promoters is sufficient to repress gene expression. The experiment compared control cells to those treated with a compound (doxycycline, or dox) that triggered the expression of an engineered zinc finger-DNA methyltransferase (ZF-DNMT) protein complex. The ZF-DNMT complex was designed to bind to and forcibly methylate the CG-rich promoter sequence of the *SOX2* gene, but in fact was found to bind nonspecifically to CG-rich sequences. The off-target effects of the system resulted in thousands of endogeneous promoters being forcibly methylated. The authors measured methylation level, gene expression, as well as active chromatin marks H3K4me3 and RNA Polymerase II (RNA PolII), to evaluate whether forcible methylation of promoters resulted in decreased signal of active transcription.

Ford et al. (2017) report that in general the expression of promoter-methylated genes in treated cells did not differ substantially compared to untreated cells. In addition, their analysis did not reveal a decrease in active chromatin state signals as a result of DNA methylation. However, our reanalysis suggests that the choice of statistical techniques and presentation of the statistical evidence hindered the ability to reliably assess the signal in the data. First, in determining which promoters were differentially methylated, Ford et al. (2017) did not make use of statistical inference. Instead, they used effect-size cutoffs without any attempt to control the rate of false discoveries arising from natural biological variability. In addition, in determining which genes were differentially expressed (DE), the normalization procedure did not properly account for the global expression differences induced by methylation. This resulted in artificial inflation of expression values in the treated cells which resulted in a reduction in the power to detect DE genes. Finally, when assessing the correlation between promoter methylation and epigenetic marks of transcription, Ford et al. (2017) used absolute differences rather than log fold changes. This choice resulted in a prioritization of relatively small differences between large counts over large-magnitude changes in small to medium counts.

Here we reassess the repressive capacity of promoter DNA methylation through an extensive analysis that overcomes the limitations outlined above. Specifically, instead of using effect-size cutoffs, we identified promoters that were significantly methylated while accurately controlling the false discovery rate using dmrseq (Korthauer et al., 2018). This prevents falsely detecting regions with large observed effect-sizes that are not statistically significant due to either very low coverage or high within-group variability. In addition, instead of global size factor normalization of RNA-Seq counts, we identify a set of control genes that are not bound by ZF-DNTMA or near a differentially methylated region (DMR) and calculate size factors from these control genes (Risso et al., 2014). By doing this we gain increased power to detect genes with differential expression due to promoter DNA methylation. Finally, when evaluating the relationship between epigenetic marks of active transcription, we consider that log fold changes, rather than absolute differences, are the more relevant signal of change (Goentoro and Kirschner, 2009).

Together, these strategies enabled us to rigorously reevaluate the causal role of forced DNA methylation of thousands of promoters. As previous work has suggested that correlations between methylation and gene repression are stronger in promoters with a CG island (Jones, 2012), we also investigate whether these associations are evident for promoters without CG islands.

## Results

### Forced promoter methylation directly induces transcriptional repression

Using the unmethylated regions (UMRs) and effect size criteria as defined by Ford et al. (2017), we replicated their figures that appear to show there is no striking relationship between promoter DNA methylation and gene expression (1A and B). However, after performing statistical inference of DMRs using dmrseq (Korthauer et al., 2018) and control-gene normalization of RNA-seq counts, a strong association is evident (Figure 1C and D). Out of all genes with significantly methylated promoters (dmrseq, FDR < 0.01), 83.6% (82.1-85.1 95% CI) have decreased expression, and 65.2% (63.3-67.1 95% CI) are significantly repressed (DESeq2, FDR < 0.10). By using dmrseq, we are able to rank DMRs by statistical significance, which presents an opportunity to assess the signal at varying levels of certainty of promoter methylation. We find that the association between promoter DNA methylation and transcriptional repression in general is stronger for more significant DMRs (Figure 2 and S1). The relationship is also slightly stronger when restricting the analysis only to genes that are expressed in control samples (Figure S2).

**Figure 1:**
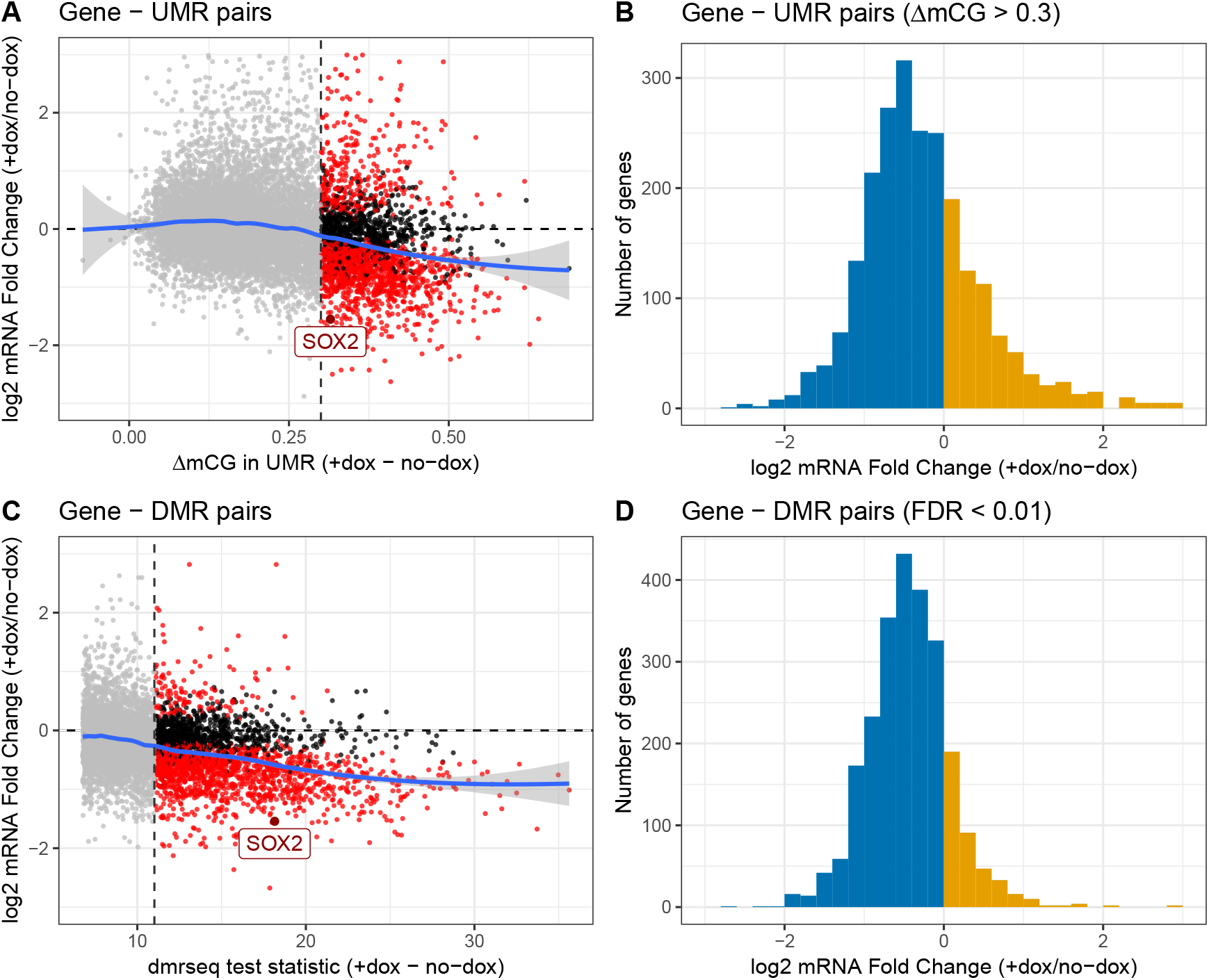
Statistical inference uncovers repressive capacity of promoter DNA methylation. (A) Analysis presented by Ford et al. (2017): Methylation difference (+dox - no-dox) in UMRs versus log2 mRNA fold change (+dox / no-dox) of genes that intersect a UMR (transcription start site +/- 2kb). Points in grey denote ΔmCG < 0.3. (B) Analysis presented by Ford et al. (2017): Histogram of log2 mRNA fold change (+dox / no-dox) of all genes +/- 2kb of the transcription start site of UMRs with ΔmCG > 0.3. (C) Reanalysis: dmrseq test statistic for DMRs (+dox - no-dox) versus log2 mRNA fold change (+dox / no-dox) of genes that intersect a DMR (promoter region +/- 2kb). All points with DMR FDR < 0.10 are shown. Points in grey denote DMRs with FDR > 0.01. (D) Reanalysis: Histogram of log2 mRNA fold change (+dox / no-dox) of genes +/- 2kb of the promoter of DMRs with FDR < 0.01. In (A) and (C), points in red denote DE genes with FDR < 0.05. In (B) and (D), bars shaded in blue denote genes repressed by DNA methylation; bars shaded yellow denote genes activated by DNA methylation. In (A) and (B), global gene normalization is used for differential expression analyses; in (C) and (D), control gene normalization is used.

**Figure 2:**
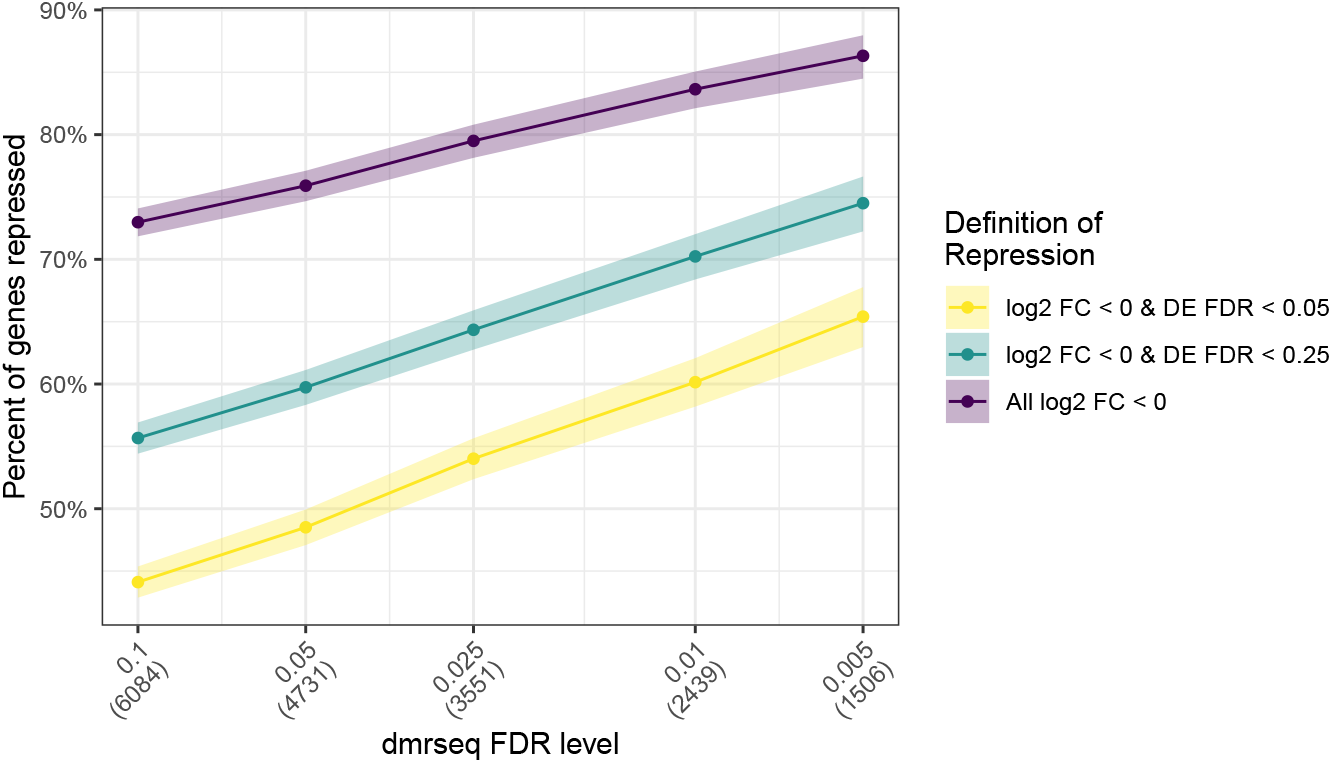
Percentage of repressed genes is higher for more significant DMRs. The percentage of significantly repressed genes (y-axis) is shown for various dmrseq FDR cutoffs (x-axis). Various DESeq2 FDR cutoffs are shown by separate lines. All genes with an average of at least 50 normalized counts in the control samples are shown.

The advantages of performing statistical inference using dmrseq are also demonstrated by comparing results to those using UMRs or DMRs identified by Ford et al. (2017). The SOX2 gene, which is the primary target of the ZF-DNMT assay, has a dmrseq statistic at the 82nd percentile of DMRs with FDR < 0.01 (Figure 1C). In contrast, the absolute methylation difference of SOX2 is only at the 14th percentile of UMRs with a difference above 0.3 (Figure 1A). Furthermore, we demonstrate that many of the DMRs found by Ford et al. (2017) using DSS (Park and Wu, 2016) exhibit substantial variability from sample to sample or contain very few CGs (Figure S3B), which could likely be false positives. In contrast, dmrseq finds many DMRs not found by DSS that exhibit clear differences between sample groups sustained over many CGs (Figure S3A)

In addition to the use of statistical inference of DMRs, this analysis also benefits from an alternative approach to RNA-Seq normalization compared to that carried out by Ford et al. (2017). Specifically, instead of standard global normalization, we compute size factors using a set of control genes that are not located near either a ZF-DNMT binding site or differentially methylated region (Methods). The distribution of control-gene normalized counts in treated samples is shifted slightly lower than the control samples (Figure S4C), demonstrating global differences between the sample groups that are not preserved under global normalization (Figure S4A and B). Under global normalization, 5,988 differentially expressed genes are detected (DESeq2, FDR < 0.05). With our approach 6,434 differentially expressed genes are identified, including all 5,988 genes detected using global normalization plus an additional 446 genes. These additional 446 genes are enriched for repression which demonstrates that control gene normalization increases power to detect significantly repressed genes (Figure S5).

### Forcibly methylated DNA shows reduced active chromatin marks

Ford et al. (2017) did not find a significant correlation between promoter DNA methylation and absolute difference in H3K4me3 read counts. However, if we represent the difference in H3K4me3 occupancy with relative fold changes rather than absolute counts (Methods), we see that promoter DNA methylation is strongly associated with reduced H3K4me3 occupancy (Figure 3A and B). In addition, we found an even stronger association using inference by dmrseq (Figure 3B) compared to absolute differences in mCG levels (Figure 3A). Out of all H3K4me3 peaks (MACS2) with significantly methylated promoters (dmrseq, FDR < 0.01), 88.6% (87.6-89.6 95% CI) have decreased H3K4me3 fold change. Although not evaluated in (Ford et al., 2017), we find a similar but slightly weaker trend for RNA PolII occupancy (Figure 3C and D). Out of all RNA PolII peaks (MACS2) with significantly methylated promoters (dmrseq, FDR < 0.01), 64.3% (63.5-66.0 95% CI) have decreased RNA PollII fold change.

**Figure 3:**
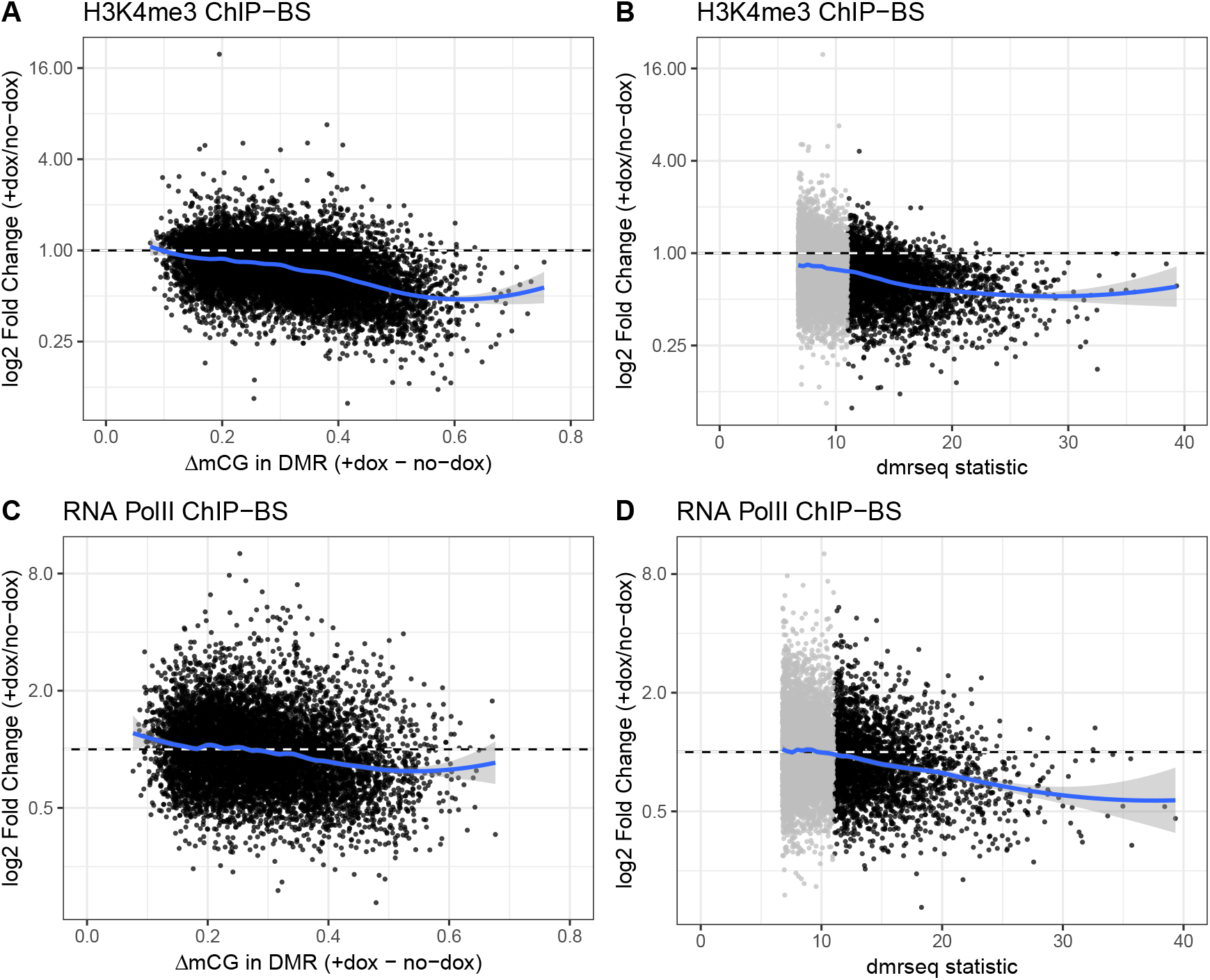
Promoter DNA methylation removes epigenetic marks of active transcription. (A) Methylation difference (+dox - no-dox) in DMRs versus log2 H3K4me4 fold change (+dox / no-dox) of peaks that intersect a DMR. (B) dmrseq test statistic for DMRs (+dox - no-dox) versus log2 H3K4me4 fold change (+dox / no-dox) of peaks that intersect a DMR. (C) Methylation difference (+dox - no-dox) in DMRs versus log2 RNA PolII fold change (+dox / no-dox) of peaks that intersect a DMR. (D) dmrseq test statistic for DMRs (+dox - no-dox) versus log2 RNA PolII fold change (+dox / no-dox) of peaks that intersect a DMR. In all panels, all points with DMR FDR < 0.10 are shown. In (B) and (D), points shaded grey denote DMRs with FDR > 0.01.

Ford et al. (2017) also concluded that forcibly methylated DNA associates with active chromatin marks, based on the increase in methylation level of UMRs of treated samples in H3K4me3 and RNA PolII peaks. We found that the same is true for dmrseq DMRs (Figure 4A and B, left panel). However, because the set of UMRs/DMRs are selected to be regions with large difference in methylation between treatment groups this is a biased comparison. Thus, we also compared these results to the differences in the remaining peaks. We find that although the methylation level in H3K4me3 and RNA PolII peaks outside of DMRs is still higher in treated samples, the difference between the treated and control is much less pronounced, especially for RNA PolII (Figure 4A and B, right panel). In addition, there is a small fraction of highly methylated peaks (mCG > 0.5) in both the control and treated samples.

**Figure 4:**
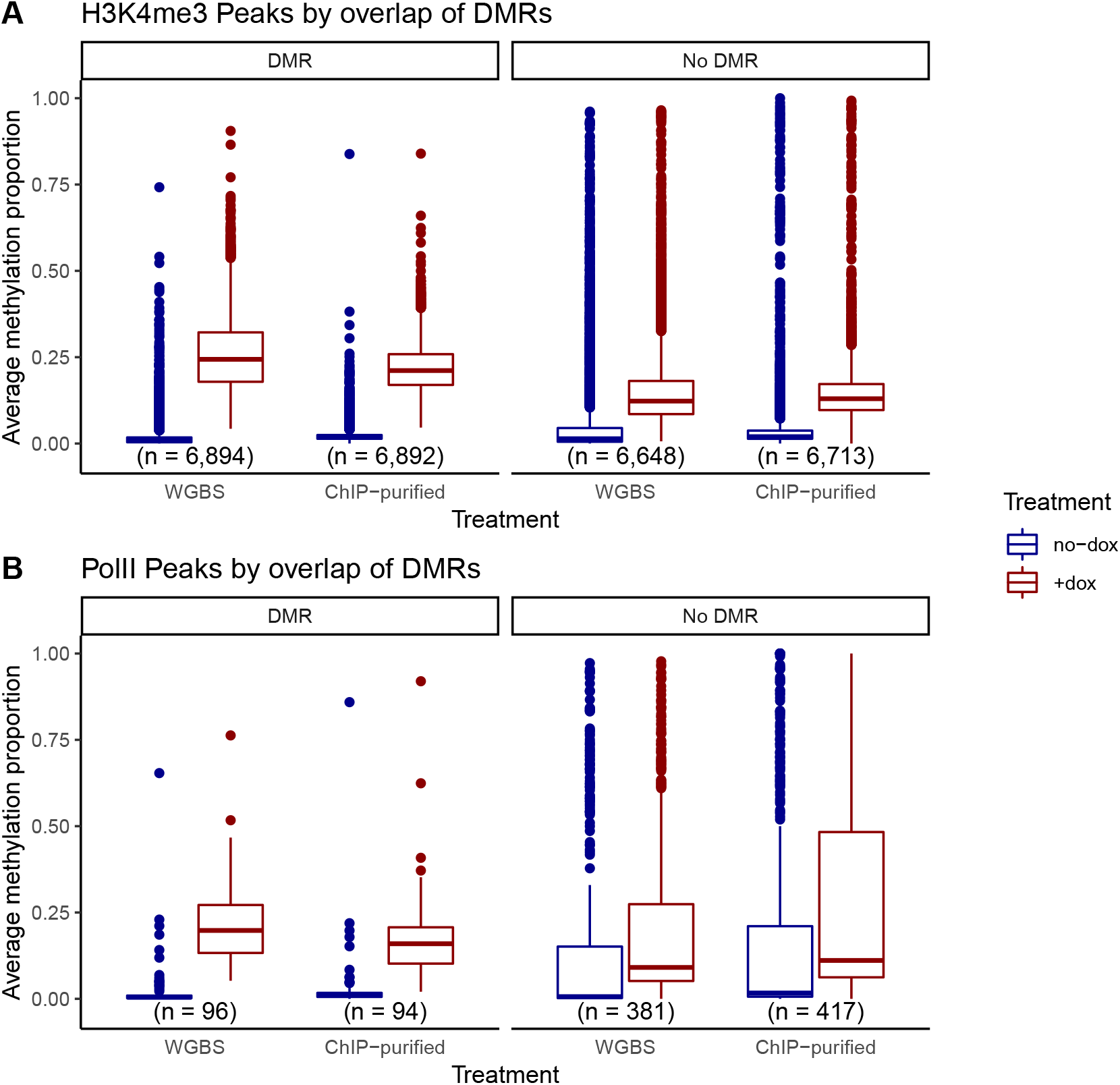
Actively transcribed regions are lowly methylated. (A) Boxplots of average methylation proportion in WGBS and ChIP-purified samples across regions that intersect H3K4me3 peaks, colored by treatment and faceted by overlap with DMRs. (B) Boxplots of average methylation proportion in WGBS and ChIP-purified samples across regions that intersect RNA PolII peaks, colored by treatment and faceted by overlap with DMRs.

### Transcriptional silencing is independent of promoter CG density

Non-CG island promoter methylation has been shown to be more transient, and the probable causal mechanism for gene silencing is less characterized than for CG island promoters (Jones, 2012; Schübeler, 2015). For this reason, it is plausible that the causal role of promoter DNA methylation may be restricted to those with a CG island. Thus, we investigated whether the strong associations we observed were sensitive to the CG density of promoters.

We find that the proportion of genes with significantly methylated promoters (dmrseq, FDR < 0.01) that are repressed (log2 fold change < 0) is not significantly different for promoters with a CG island and those without (Figure 5, *χ*^2^ p = 0.13). We also find that the pattern of increased methylation and decreased expression is similar for both types of promoters (Figures S6 and S7). Likewise, there is no evidence for a substantially different trend in the pattern of increased methylation with decreased H3K4me3 and RNA PolII occupancy (Figure S8).

**Figure 5:**
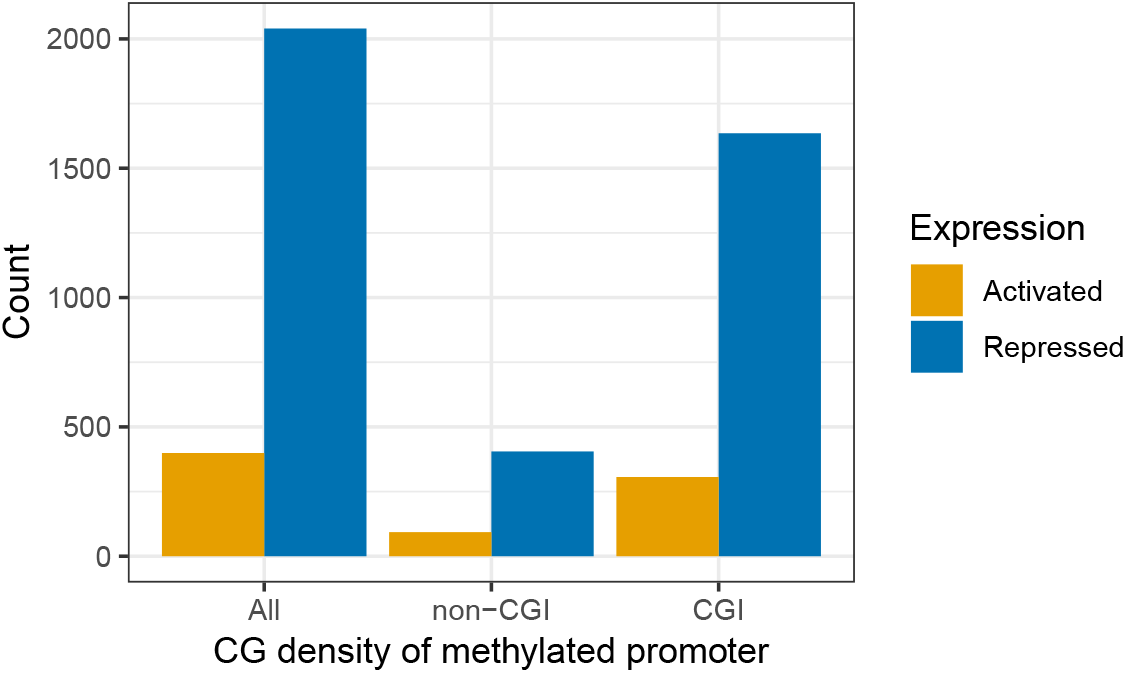
CG-poor promoters show similar repressive capacity to CG islands. Barplot of the number of genes with significantly differentially methylated promoters (FDR < 0.01, +/- 2kb) that are repressed (log2 fold change < 0) or activated (log2 fold change > 0). Results are shown for all promoters, promoters that do not overlap a CG island (non-CGI) and promoters that overlap a CG island (CGI), and color denotes direction of effect.

## Discussion

Thanks to the commitment to open science by Ford et al. (2017), we have conducted a thorough reassessment of whether forced promoter DNA methylation is generally sufficient for transcriptional repression. With the use of an approach that provides statistical inference for DMR detection, a normalization technique capable of retaining global differences, and using log fold change to quantify differences in count data rather than absolute differences, we have come to an opposite conclusions than the original study. Specifically, we have demonstrated that promoter methylation directly represses gene expression in the vast majority of differentially expressed genes, and reduces the level of complementary epigenetic markers of active transcription.

We note that although the strength of association between forced promoter DNA methylation and decreased H3K4me3 and RNA PolII occupancy was not as strong as that of gene expression, this is somewhat expected since biological replicates were available for the latter but not the former. With only one replicate per condition in the ChIP-BS data, we cannot be certain that the peaks we obtain are statistically significant and thus the association analysis will unavoidably include higher level of noise. This may also explain the higher methylation level of H3K4me3 peaks in treated samples, as well as the apparent coexistence of methylation and active chromatin marks in control samples. More investigation is needed to evaluate whether these patterns hold for significant peaks called from replicate samples.

We also note that although the effect of forced promoter DNA methylation on transcription is clear and strong, this effect is observed in the samples treated for three days, with no withdrawal period. As noted in the original study, the induced methylation is not permanent and after a withdrawal period of nine days the methylation levels are regressed closer to the levels at baseline. Thus, although this study provides evidence for a causal role in the initiation of transcriptional silencing, it suggests that other factors are necessary for its continued maintenance. Further work on the stability and dynamics of transcriptional repression is warranted.

Details of our analysis are included below and the code needed to reproduce our findings is included in a public GitHub repository at github.com/kdkorthauer/repressivecapacity.

## Methods

Note that all references to Ford et al. (2017) refer to the current version of the preprint at the time of this writing (the version posted on September 20, 2017). All analyses were carried out using R (R Core Team, 2016) version 3.5.0 unless otherwise specified.

### Data Download

Supplementary data tables from Ford et al. (2017): Supplementary tables S1-S4 (files 170506-1.txt-170506-4.txt, respectively) were downloaded from the bioRxiv website at https://www.biorxiv.org/highwire/filestream/57972/field_highwire_adjunct_files/.

RNA-Seq read counts: The file GSE102395_MCF7_ZF_DNMT3A_countstable.txt was downloaded from GEO accession number GSE102395. The eight control samples (untreated with dox) and four treated ZF-DNMT samples were used for the differential expression analyses.

WGBS read counts: All methylated and total coverage read count tables were downloaded from GEO accession number GSE102395 using the R/Bioconductor package GEOquery (Davis and Meltzer, 2007), with the exception of one file which was found to be corrupt. The file MCF7_ZF_DNMT3A_doxInduced_rep2.allC.gz was instead downloaded from the lab website of the authors of Ford et al. (2017) at http://cpebrazor.ivec.org/public/listerlab/ethan/Public/. This was made available following personal communication with the authors regarding the corrupt file on GEO. The three control samples (empty vector and no dox ZF-DNMT) and three treated samples (ZF-DNMT dox treated) were used in the differential methylation analyses.

Raw ChIP-BS reads: fastq files for H3K4me3 and RNA PolII treatment and control samples (1 replicate each) were downloaded from GEO accession number GSE102395 using GEOquery and the SRA Toolkit (Leinonen et al., 2011).

Annotations: Promoter and CG island annotations for hg19 were downloaded using the R/ Bioconductor package annotatr (Cavalcante and Sartor, 2017). The grch37 assembly file Homo_sapiens. GRCh37.dna.primary_assembly.fa.gz was downloaded from Ensembl release 92.

### Normalization of RNA-Seq count data

Normalization methods such as the global size factor parameter estimation in DESeq2 (Love et al., 2014) are violated in the presence of global expression differences (Risso et al., 2014). In this experiment, the ZF-DNMT compound nonspecifically binds to over 25,000 CG-rich sites, a large proportion of which overlap promoter regions (Ford et al., 2017). As such, we expect to see global expression differences if the DNA methylation has a repressive effect. In this case, without accounting for the global differences, standard global normalization will artificially inflate the counts of genes in the treated condition. This will lead to a loss of power to detect differentially expressed genes. Instead, normalization should only be done using a set of control genes, for which we do not expect differential expression. This same issue is a concern in the normalization of ChIP-BS reads, which is described in the next section.

For these reasons, we compute size factors on a set of control genes. Here we define the control genes as those that were at least 10kb away from a ZF-DNMT binding site or putative DMR (by dmrseq, with FDR level 0.25). ZF-DMNT binding sites were identified using the ChIP-seq binding site data from the Supplementary table S1 of Ford et al. (2017). There were 8009 such genes that met our criteria for control genes. We also filter out very lowly expressed genes (total counts in all samples together fewer than 10 or counts of zero in more than half of the samples).

### Preprocessing and normalization of ChIP-BS reads

Quality control of fastq files was assessed using fastqc (Andrews, 2010), and reads were quality- and adapter-trimmed using TrimGalore! (Krueger, 2015) using default settings. Bisulfite-aware mapping and cytosine methylation count extraction was performed with Bismark (Krueger and Andrews, 2011) version 0.17.0 and the GRCh37 genome assembly with the default options. Bam files were sorted with SAMtools (Li et al., 2009).

Peaks were called using MACS2 (Zhang, 2008) using the genome size option -g 2.7e9. To compute size factors, we count reads mapping to the intersection of common peaks (in treated and control samples). Following Ford et al. (2017), we restrict this calculation to peaks where both treatment and control samples have an average methylation level (using only sites with at least 5 reads total in each treatment) less than 0.2, as well as a mean difference between treatment and control of less than 0.2. We also exclude regions within 5kb of a ZF binding site, for reasons described in the previous section. This calculation is done separately for the H3K4me3 and RNA PolII samples. For downstream analyses, read counts are scaled by the size factor from this total.

### Association analyses of UMRs

The association between methylation in UMRs and gene expression is evaluated using Supplementary Table S4 from Ford et al. (2017), which contains the difference in methylation of UMRs between controls and treated samples, as well as the log2 fold change of expression in the associated genes. These values are used to generate Figure 1A and B.

### Association analysis of RNA-Seq and WGBS

Differential expression analysis comparing control to treated samples was carried out with DESeq2 (Love et al., 2014). Methylation counts were read into R using the bsseq R/Bioconductor package (Hansen et al., 2012). 257,012 CG loci (1%) with zero coverage in at least one condition were removed. Differential methylation analysis comparing control to treated samples was carried out with dmrseq (Korthauer et al., 2018) with parameter options bpSpan = 5 0 0, maxGap = 5 0 0, and maxPerms = 2 0.

DMRs that were within +/- 2kb of the promoter region of any gene were matched to the closest such gene. If the promoter of that gene contained at least 100 basepairs of a CG island, it was classified as a CG island promoter. Percentages of genes with repressed expression were calculated at various cutoffs of dmrseq (ranging from 0.005 to 0.10), and for several definitions of repression. Repression was defined as a negative log2 fold change, along with differential expression with FDR cutoffs for DESeq2 ranging from 0.05 to 1. Associations were also examined for sets of genes excluding those unexpressed in control samples, with cutoffs for expressed genes of mean normalized counts varying from 50 to 250. The number of genes meeting each cutoff is shown in Figure S9. Unless otherwise specified, all results are presented using expressed genes with a cutoff of 50. Confidence intervals for percentages (proportions) were obtained using the ‘wilson’ option of the ‘binom.confint’ function of the ‘binom’ R package (Dorai-Raj, 2014). A *χ*^2^ test was carried out to test the null hypothesis of independence of CG island status and transcriptional repression for differentially expressed genes with FDR < 0.05 and DMRs with FDR < 0.01.

### Association analyses of RNA-Seq and ChIP-BS

Mean methylation difference in DMRs were calculated from WGBS counts, over all loci that had nonzero coverage in each treatment group. If the DMR contained at least 100 basepairs of a CG island, it was classified as a CG island DMR.

The following was carried out separately for H3K4me3 and RNA PolII samples. The number of reads in each 5kb window centered on the set of DMRs (FDR < 0.10) were counted using the R/Bioconductor package GenomicAlignments (Lawrence et al., 2013), and these counts were normalized by the size factors described above. Only regions with at least 50 total normalized counts total (treated and control samples summed together). Percentages of genes with decreased normalized ChIP-BS counts were calculated at various cutoffs of dmrseq (ranging from 0.005 to 0.10), and confidence intervals were obtained using the ‘wilson’ option of the ‘binom.confint’ function of the ‘binom’ R package (Dorai-Raj, 2014).

### Comparison of ChIP-BS and WGBS

Methylation counts for WGBS and ChIP-BS were read into R using the bsseq R/Bioconductor package. Average methylation levels for common peaks (between control and treated) were calculated separately for WGBS and ChIP-BS data. Only loci with at least 3 reads per sample in the ChIP-BS data, and at least 4 reads per treatment group in the WGBS data were used for the average methylation calculation. Peaks were checked for overlap with any DMR and boxplots of average methylation levels in both experiment types were plotted separately for those that did and did not overlap a DMR.

### Data Access

All code to generate the results in this manuscript, including data download, preprocessing, analysis, and generation of figures, is included in the public GitHub repository at github.com/kdkorthauer/repressivecapacity.

## Acknowledgments

We thank Ryan Lister for valuable comments and discussion that helped to improve this manuscript. We also gratefully acknowledge Ethan Edward Ford, Matthew R. Grimmer, Sabine Stolzenburg, Ozren Bogdanovic, Alex de Mendoza, Peggy J. Farnham, Pilar Blancafort, and Ryan Lister for making the preprint and experimental data from their recent study publicly available. Without their commitment to open science and rapid dissemination of results, this work would not be possible. This work was supported by R01HG005220 from the NIH (to RAI).

---

**Figure S1.**
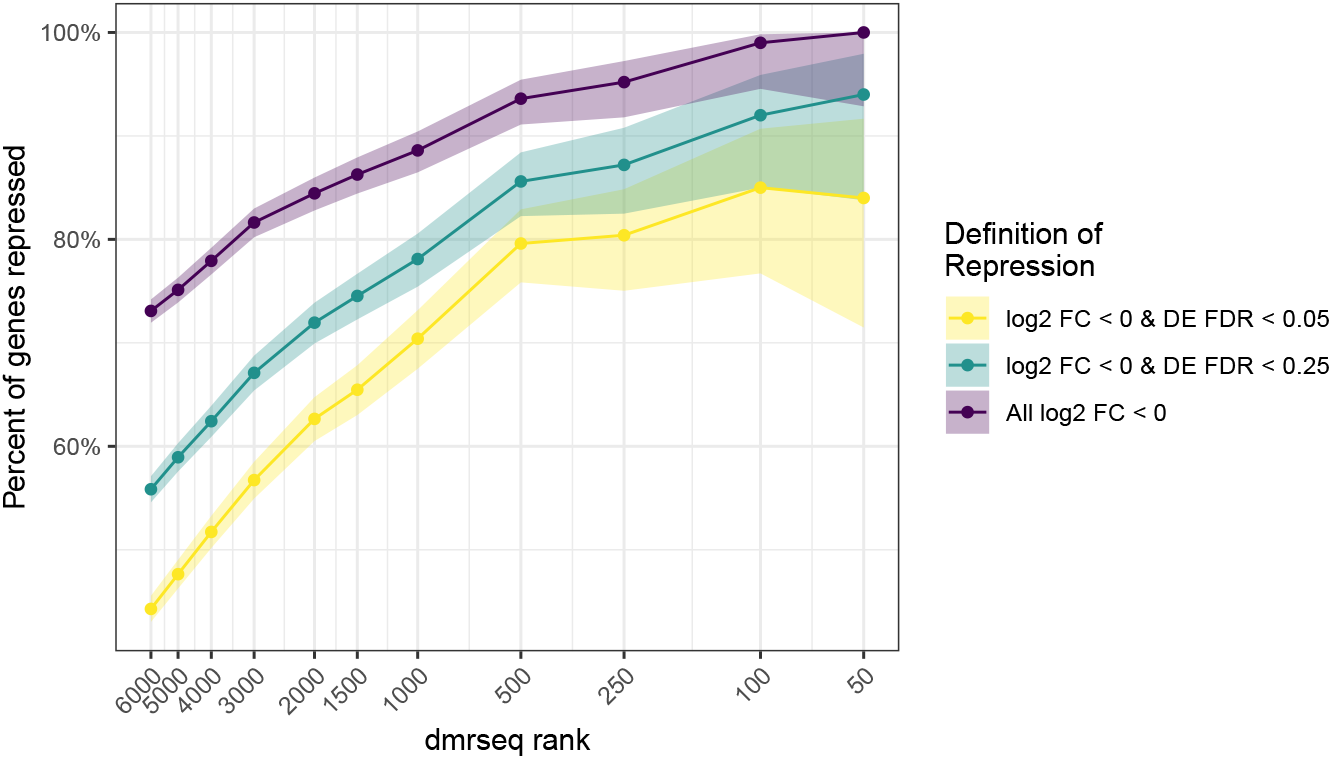
Percentage of repressed genes increases for highly ranked DMRs. The percentage of significantly repressed genes (y-axis) is shown for various dmrseq statistic cutoffs (x-axis). Various DESeq2 FDR cutoffs are shown by separate lines. All genes with an average of at least 50 normalized counts in the control samples are shown.

**Figure S2.**
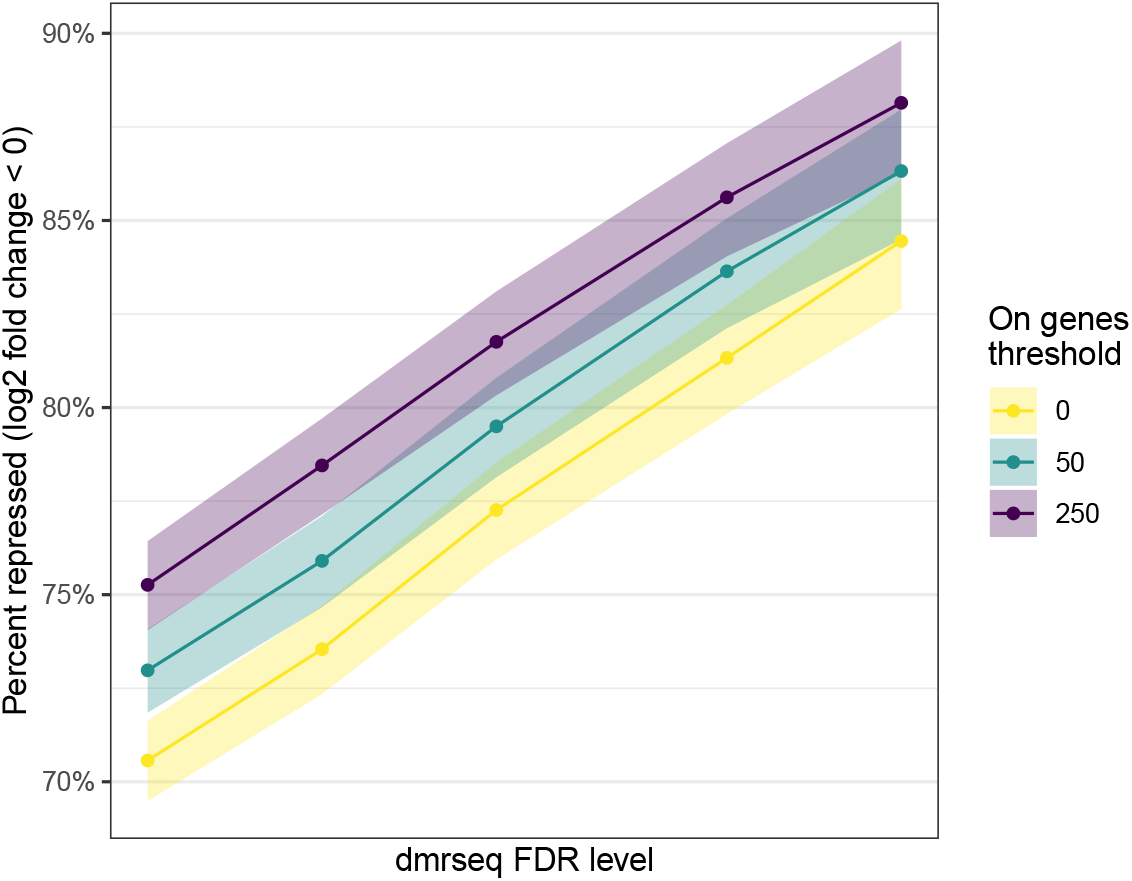
Relationship between methylation and repression is stronger for genes expressed in control samples. The percent of repressed genes (log2 fold change < 0, y-axis) is shown for various dmrseq FDR cutoffs (x-axis). The three different lines represent different thresholds of mean normalized counts of control samples (representing the genes that are ‘on’).

**Figure S3.**
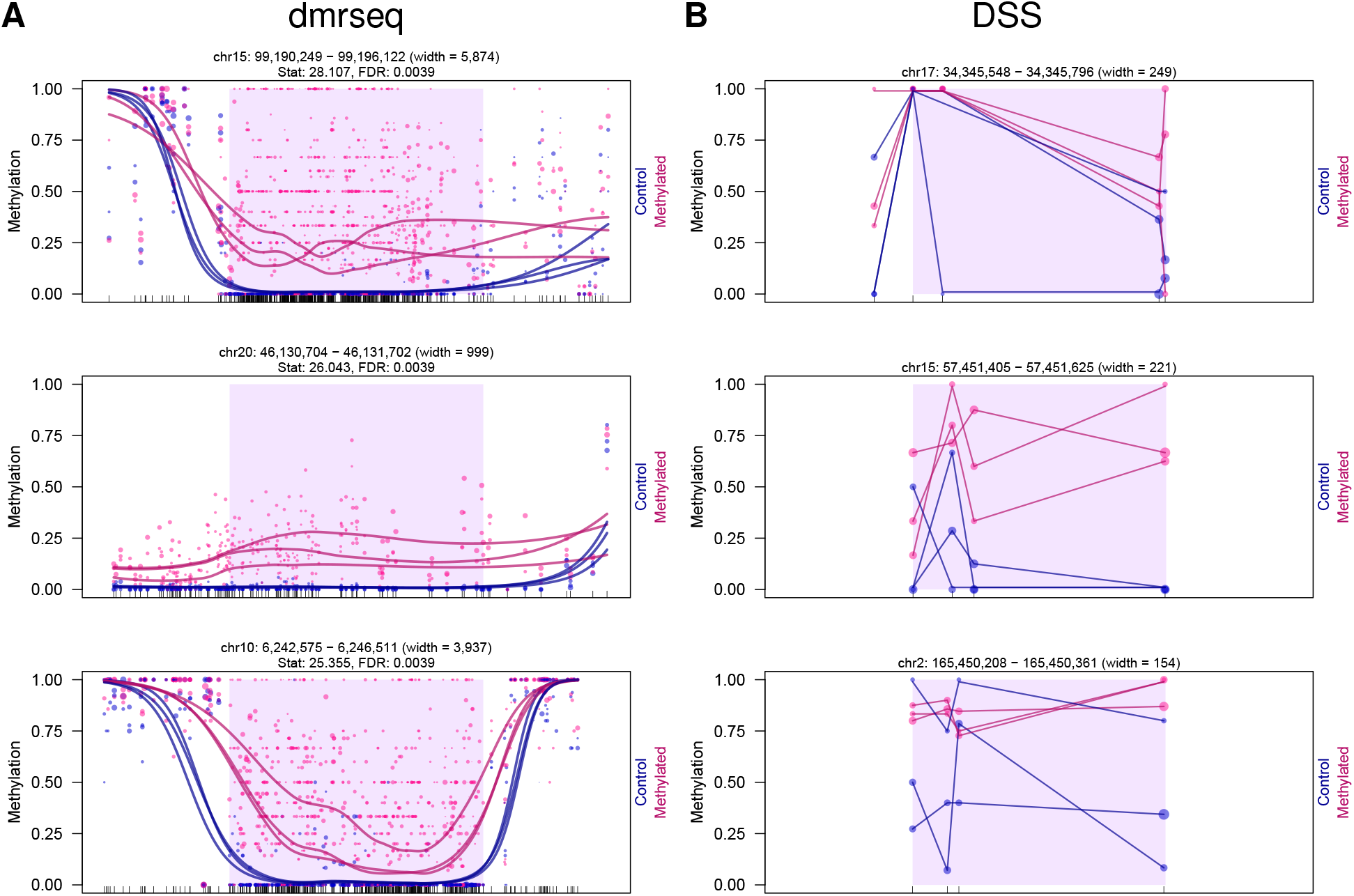
Top-ranked regions found only by dmrseq compared to those found only by DSS. The top three regions ranked by dmrseq statistic that are not found by DSS (A), and the top three regions ranked by absolute mean mCG difference that are not found by dmrseq (FDR < 0.10) (B).

**Figure S4.**
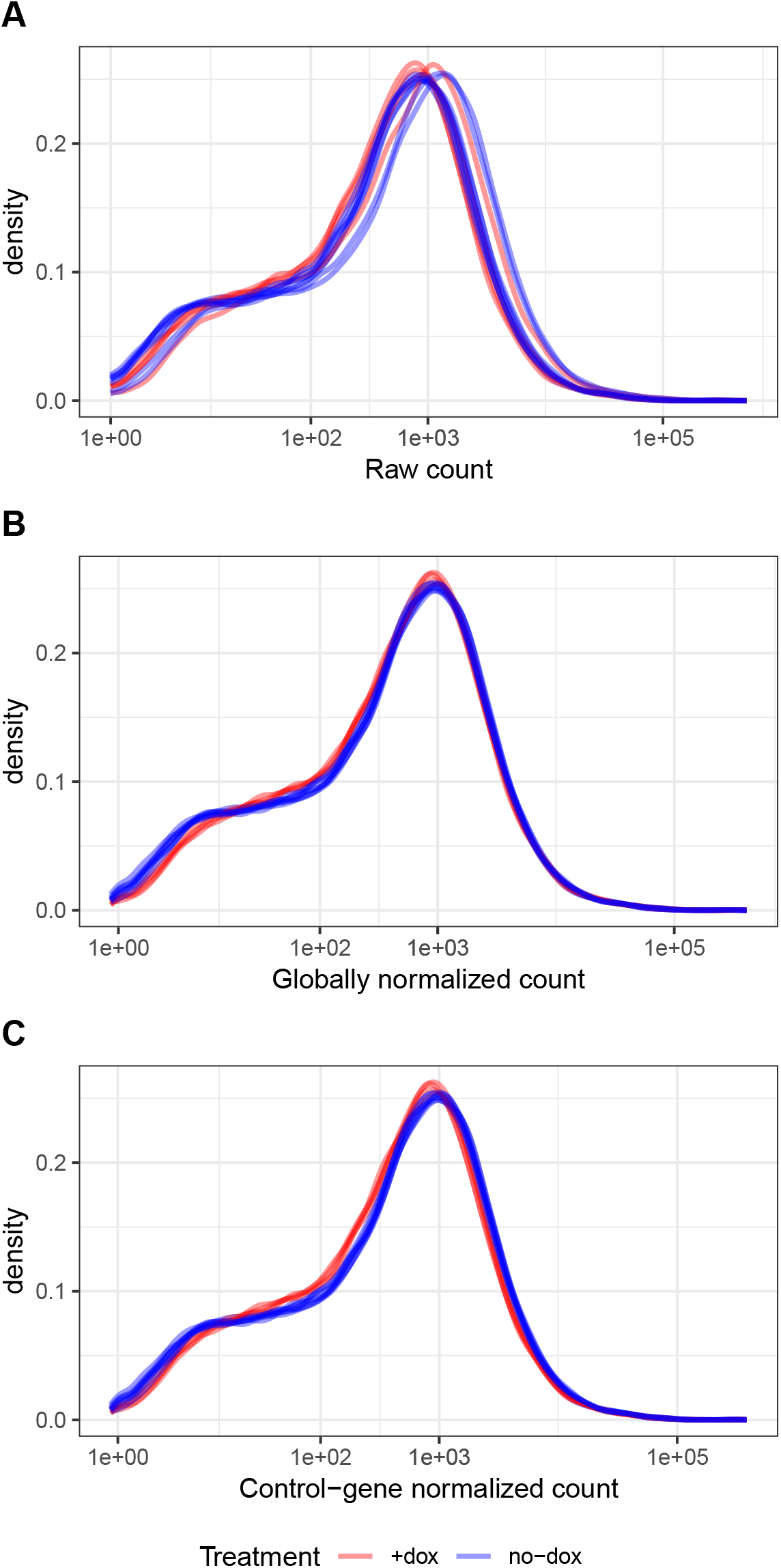
Control gene normalization reveals global expression differences. Smoothed density of (A) raw RNA-seq read counts, (B) global size factor-normalized RNA-seq read counts, and (C) control-gene size factor-normalized RNA-seq read counts.

**Figure S5.**
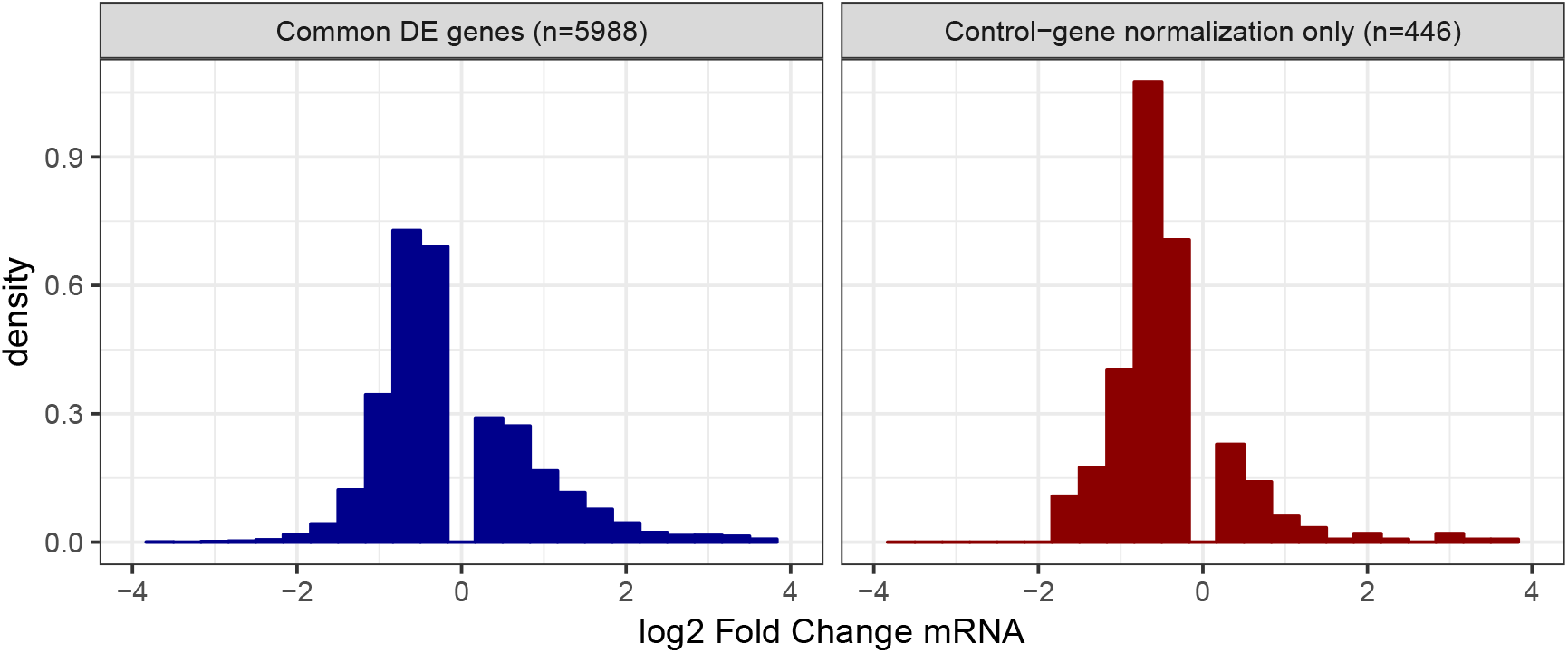
Control gene normalization increases power to detect repressed genes. Density histograms of log2 fold change (DESeq2) for all differentially expressed genes found by both global and control gene normalization (left panel, blue), as well as those genes found only by control gene normalization (right panel, red).

**Figure S6.**
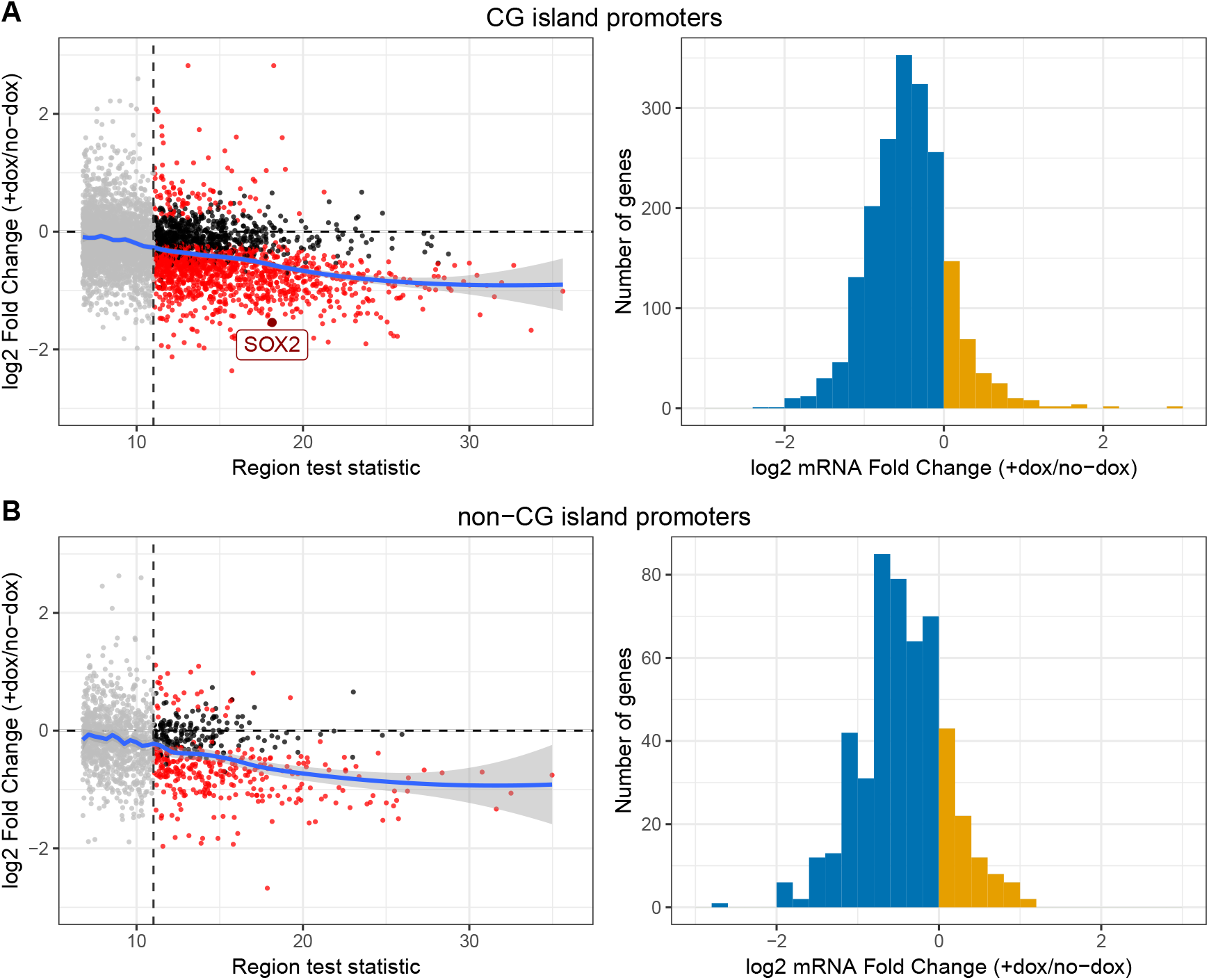
Repressive patterns are similar for CG island and non-CG island promoters. (A) dmrseq test statistic for DMRs (+dox - no-dox) versus log2 mRNA fold change (+dox / no-dox) of genes with a CG island promoter that intersect a DMR (promoter region +/- 2kb). (B) Histogram of log2 mRNA fold change (+dox / no-dox) of differentially expressed genes (FDR < 0.05) with a CG island promoter +/- 2kb of the promoter of DMRs with FDR < 0.01. (C) Same as A, but for genes without a CG island promoter. (D) Same as B, but for genes without a CG island promoter. In (A) and (B), all points with DMR FDR < 0.10 are shown. Points in grey denote DMRs with FDR > 0.01. Points in red denote differentially expressed genes with FDR < 0.05. Bars shaded in blue denote genes repressed by DNA methylation; bars shaded yellow denote genes activated by DNA methylation.

**Figure S7.**
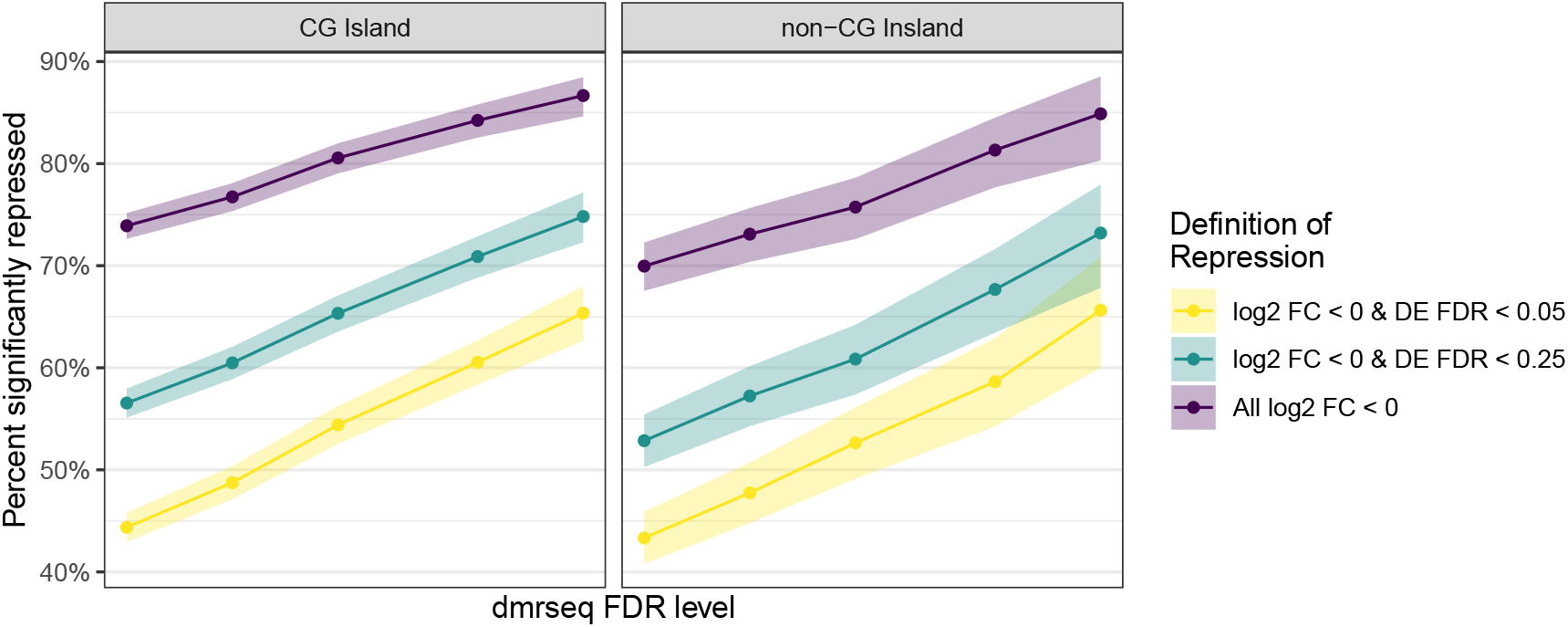
Relationship between methylation and repression is similar for CG island and non-CG island promoters. The probability of significant gene repression (y-axis) is shown for various dmrseq FDR cutoffs (x-axis). Various DESeq2 FDR cutoffs are shown by separate lines. All genes with an average of at least 50 normalized counts in the control samples are shown. Results are faceted by genes with a CG island promoter (CGI) and those without (non-CGI).

**Figure S8:**
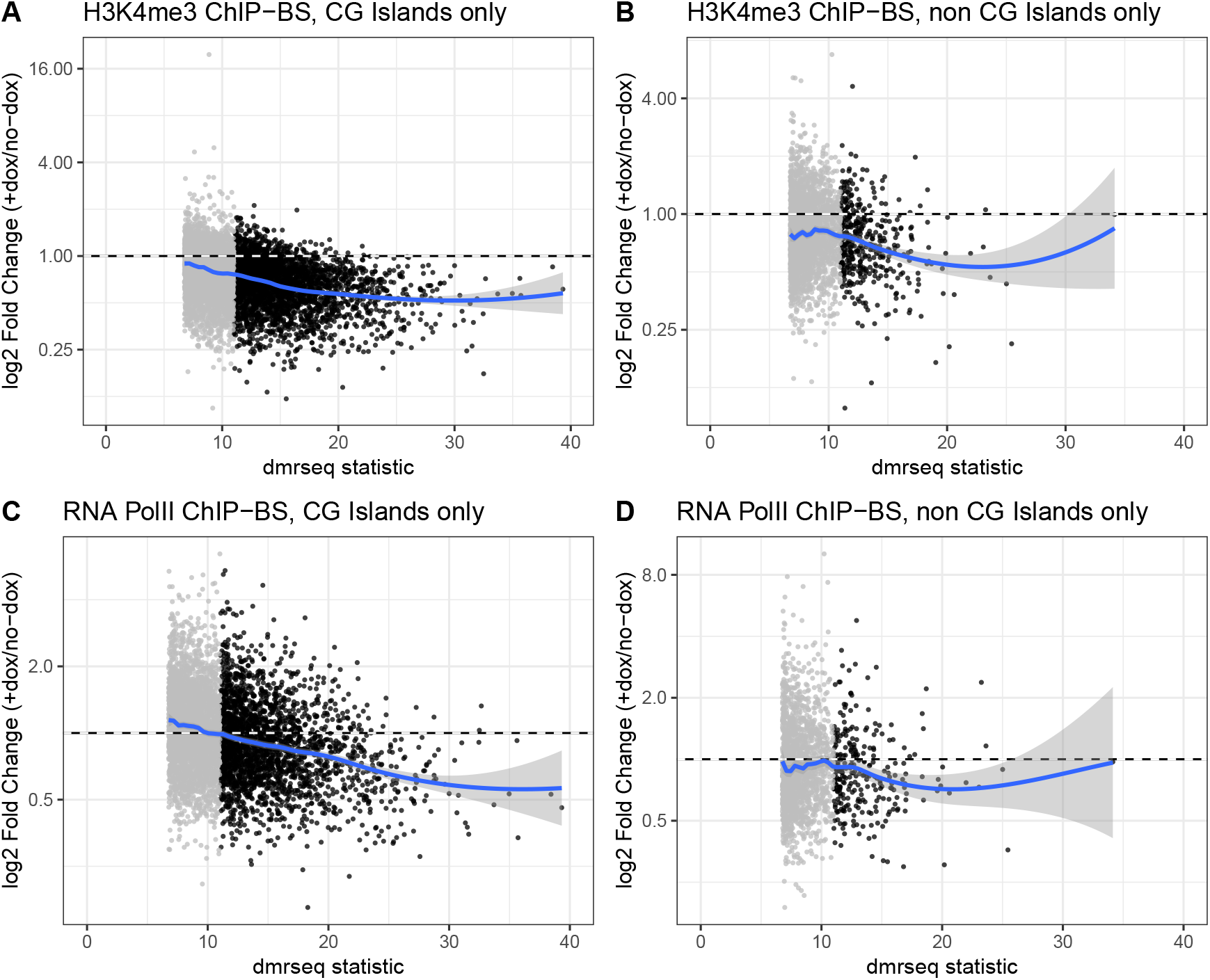
Reduction of active transcription markers is similar in CG island and non-CG island. (A) dmrseq test statistic for DMRs (+dox - no-dox) versus log2 H3K4me4 fold change (+dox / no-dox) of CG island peaks that intersect a DMR. (B) Same as (A) but for peaks that do not overlap a CG island. (C) dmrseq test statistic for DMRs (+dox - no-dox) versus log2 RNA PolII fold change (+dox / no-dox) of peaks that intersect a DMR. (D) Same as (C) but for peaks that do not overlap a CG island. In each panel, all points with DMR FDR < 0.10 are shown, and points shaded grey denote DMRs with FDR > 0.01. g(A)

**Figure S9.**
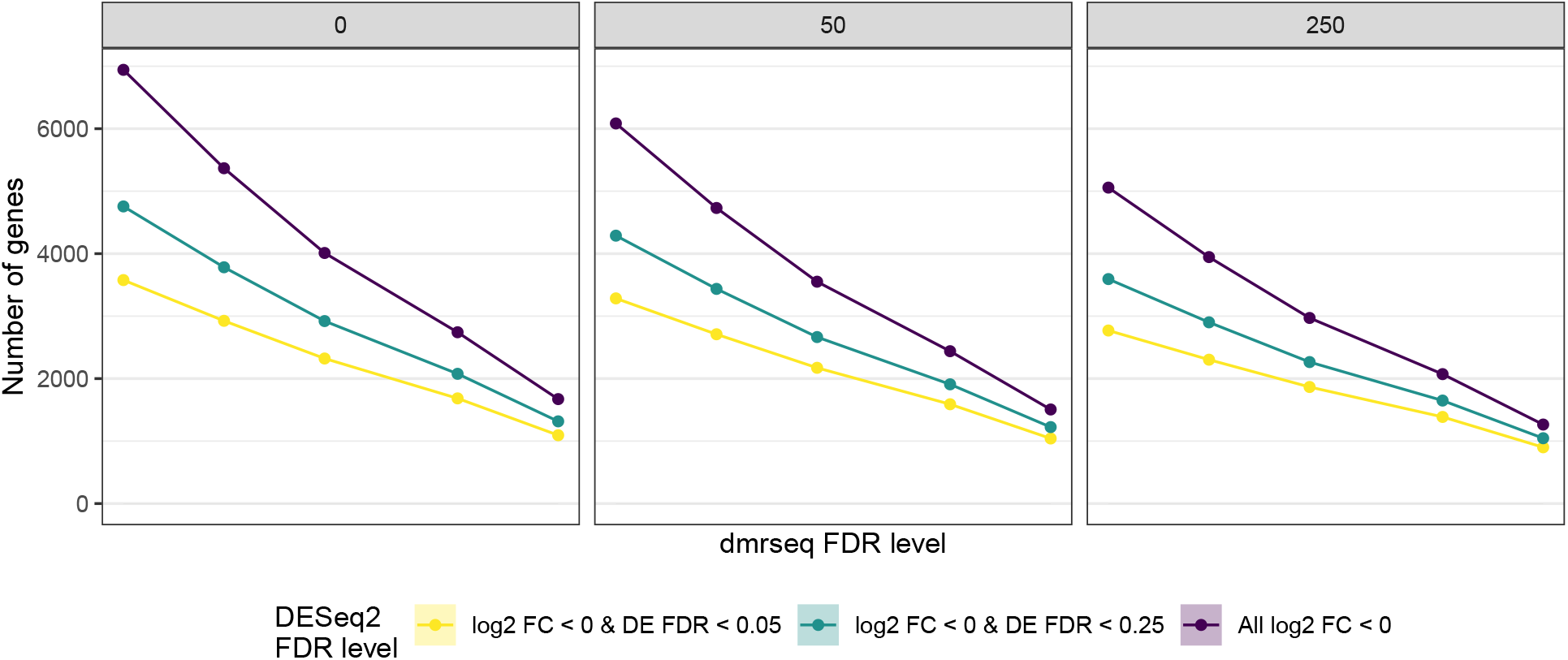
Number of genes at each FDR cutoff. The number of genes passing the cutoffs (y-axis) is shown for various dmrseq FDR levels (x-axis). Various DESeq2 FDR cutoffs are shown by separate lines. Results are faceted by various thresholds of mean normalized counts of control samples. In each panel, all genes with mean normalized count above the given threshold are included.

